# Genome of the North American wild apple species *Malus angustifolia*

**DOI:** 10.1101/2023.11.16.567428

**Authors:** Ben N. Mansfeld, Shujun Ou, Erik Burchard, Alan Yocca, Alex Harkess, Ben Gutierrez, Steve van Nocker, Lisa Tang, Christopher Gottschalk

**Affiliations:** Department of Biology, Washington University in St. Louis, St. Louis, Mo; Department of Molecular Genetics, The Ohio State University, Columbus, OH; USDA ARS, Appalachian Fruit Research Station, Kearneysville, WV; HudsonAlpha Institute for Biotechnology, Huntsville, AL; USDA ARS, Plant Genetic Resource Unit, Geneva, NY; Department of Horticulture, Michigan State University, East Lansing, MI

## Abstract

Apple (*Malus* × *domestica* Borkh.) production faces many challenges stemming from abiotic and biotic stresses. Abiotic stressors, such as extreme temperatures, droughts, and spring frosts, can lead to diminished yields and tree loss, while biotic stresses like fire blight and pest infestations further reduce tree health and fruit quality. To lessen the threat of these challenges, plant breeders aim to introduce resistance and resilience genes into cultivated varieties. However, high-relatedness among cultivated varieties and breeding lines, coupled with the long juvenility and generation times in apples, hinder the breeding process. The introduction of resistance traits from wild relatives is also constrained by these factors, as well as the lack of genomic resources that could assist in accelerating the introgression process. Herein, we report the assembly and annotation of *Malus angustifolia*, the Southern Crabapple, one of Eastern North America’s native species. Using a combination of Pacific Biosciences High Fidelity reads, Next-generation short read sequencing, as well as chromatin conformation capture sequencing, we achieve an extremely contiguous haplotype-resolved assembly. We perform comparative haplotypic analyses to identify SNPs and large structural variants, shedding light on the genomic landscape of *M. angustifolia*. Finally, we explore the phylogenetic and syntenic relationships between Eurasian *Malus* progenitors and the recently sequenced North American species, contributing valuable insights to the broader understanding of apple evolution and potential breeding strategies.

## Introduction

Apple (*Malus* × *domestica* Borkh) production faces numerous challenges arising from both abiotic and biotic stresses (Volk et al., 2015a). Abiotic stressors, such as high temperatures, droughts, and spring frosts, can result in reduced yields, stunted growth, and, in extreme cases, the loss of entire trees (Atkinson et al., 2000; Bhusal et al., 2019; Dalhaus et al., 2020; Schrader et al., 2001; Torres et al., 2013, 2016). Biotic stresses including devastating diseases such as fire blight, and pest infestations, can further diminish yields or render the produced fruit unmarketable (MacHardy, 1996; Chatterjee, 2001). To overcome these limitations in apple production, plant breeders aim to introduce resistance or resilience genes into improved cultivars. However, in the case of apples, this process has been hindered by breeding bottlenecks resulting from the high degree of relatedness among many cultivated varieties and breeding lines (Migicovsky et al., 2021; Muranty et al., 2020). Introgression breeding from wild relatives in apple is rare due to long juvenility and generation times (Fischer 2004). These challenges further amplify the time required to breed out unwanted traits and break the linkage drag associated with introgression breeding using wild crop relatives (Crosby et al., 1992).

*Malus angustifolia*, commonly known as the Southern Crabapple, is one of four native North American species (Volk et al., 2015a). With a natural range spanning from Texas to Pennsylvania, *Malus angustifolia* thrives in a variety of ecological niches including valleys and lowlands to the borders of woodlands and agricultural fields (USDA NRCS, 2002). *M. angustifolia* is typically found in areas where the soil is well drained, moist and acidic (USDA NRCS, 2002). This wild apple species frequently faces conditions that would prove intolerable to the vast majority of *M.* × *domestica* cultivars. For example, *M. angustifolia* is an extremely late blooming species (Gottschalk & van Nocker, 2013) and grows in regions affected by high temperatures, short dormancy season, and severe weather conditions (Konrad II & Fuhrmann, 2013). Additionally, like its other North American counterparts*, M. angustifolia* has co-evolved with *Erwinia amylova*, the devastating bacteria that is responsible for the fire blight disease is endemic to North America (Bonn & van der Zwet, 2000; Chatterjee, 2001). Resistance to fire blight has been observed by most of the USDA accessions (Dougherty et al., 2021; Khan & Chao, 2017; Emeriewen et al., 2014). Our recent exploration into the Pacific Crabapple, *Malus fusca,* genome suggests that this co-evolution in North America likely manifested as presence and absence variation of G-type receptor like genes, at a known resistance locus for fire blight, which may contribute to resistance to this disease (Mansfeld et al., 2023). Thus, the North American wild *Malus* relatives are poised to act as a crucial and valuable trove of genetics to improve apple resistance to biotic and resilience abiotic threats.

To address some of these challenges related to the integration of wild apple genes into apple cultivation, considerable efforts have been directed toward sequencing and assembling wild *Malus* genomes and developing other genomic tools that can facilitate the use of wild *Malus* species in cultivar enhancement (Mansfeld et al., 2023; Ruhsam et al., 2022; Li et al., 2022; Sun et al., 2020; Chen et al., 2019). As part of our effort to continue to explore and develop genomes of North American *Malus* species, we herein report the high-quality diploid genome assembly and annotation of *M. angustifolia* accession PI 613880. *M. angustifolia* PI 613880 is cataloged as “very resistant/no occurrence” to fire blight and is “extremely late blooming” (USDA Agriculture Research Service, 2015; Gottschalk & van Nocker 2013), desirable traits to target for breeding of frost avoidance and fire blight resistant apples. This accession was collected in Berkeley Co., near Holly Hill, South Carolina in 1987 by Elizabeth Dickson of the NYS Agriculture Experimental Research Station (*i.e.*, Cornell AgriTech) of Cornell University (Fig. 1A and B; USDA Agriculture Research Service, 2015). To this end we employed a mixture of Pacific Biosciences (PacBio) High Fidelity (HiFi) long reads, short Illumina reads as well as high-throughput chromosome conformation capture (Hi-C) combined with the latest genome assembly and annotation software to generate a reference quality genome for *M. angustifolia*.

**Figure 1.**
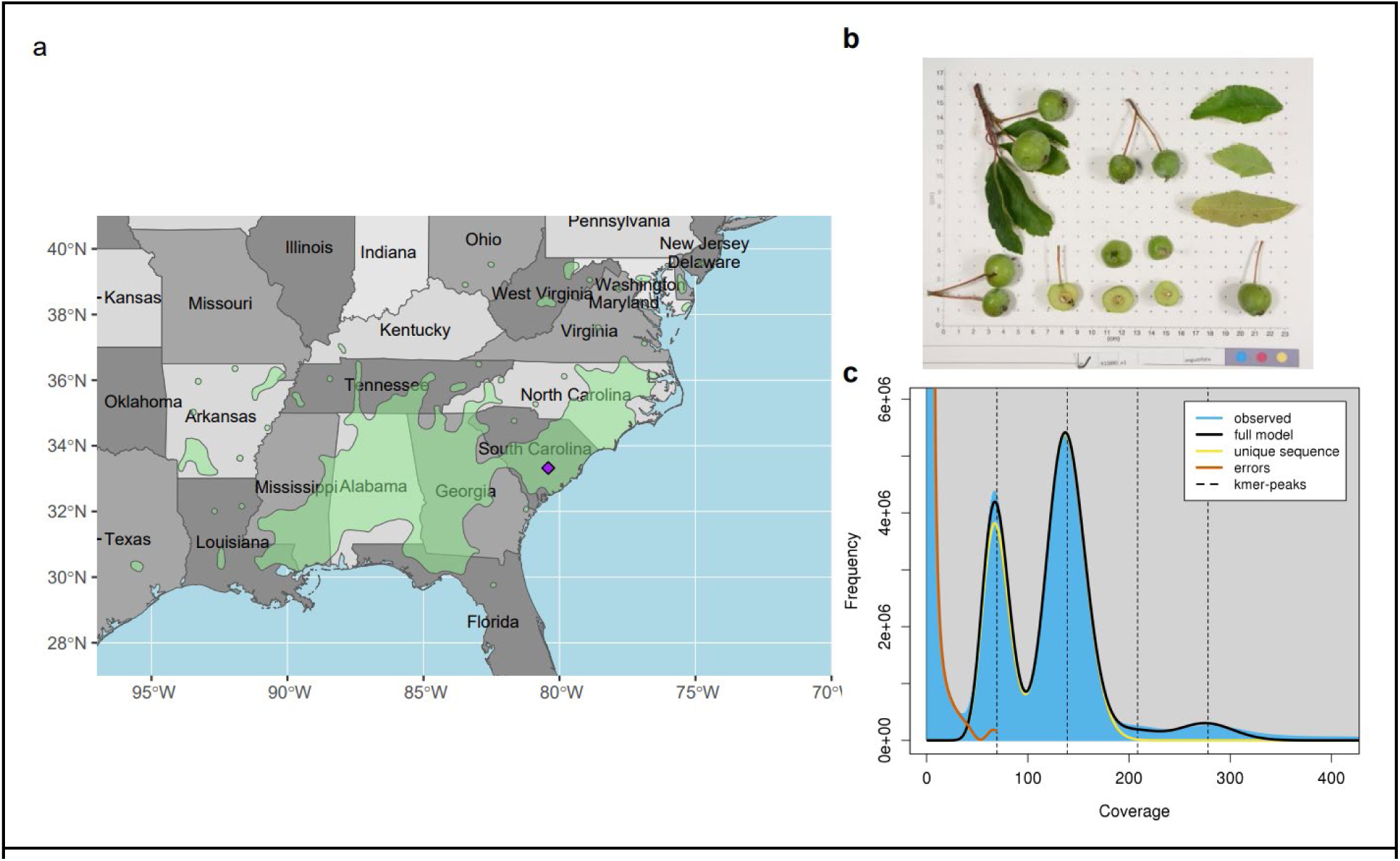
A) Map of the approximate collection position of the *Malus angustifolia* PI 613880 in South Carolina. Insert is of the specific location in Berkeley Co. near Holly Hill, SC.. B) *Malus angustifolia* accession PI 613880 GRIN-Global identification photograph. C) *k*-mer count plot from GenomeScope. The haploid genome size was predicted to be 747 Mb with approximately 58% non-unique sequence (*i.e.,* repeats) and an estimated heterozygosity of 1.06%.

## Methods

### Plant Materials

Dormant scion cuttings of PI 613880 were obtained from the USDA *Malus* Collection in Geneva, NY in February 2021. The scions were placed into a beaker with a rehydration solution (Rose 100, Floralife, Waterboro, SC) using the manufacturer’s recommended concentration and placed under long-day conditions at 21°C in a growth chamber to force bud break. Once leaves on the scions reached 3 cm in length, the scions were transferred to dark conditions for 48 hours at 21°C. Leaves were collected, weighed and split into 1 g samples. The tissue samples were placed into 5 mL centrifuge tubes and flash frozen in liquid N_2_ and stored at −80°C for later use. Additional leaf tissue was collected from living trees in the USDA *Malus* Collection in May and June of 2021, and shipped overnight in plastic bags with a damp paper towel. Upon receipt, leaves were weighed and split into 0.25 g samples. These samples were placed into 5 mL centrifuge tubes and flash frozen in liquid N_2_ and stored at −80°C. The GPS coordinates of the sample collection was retrieved from the Germplasm Resource Information Network (GRIN) passport info (https://npgsweb.ars-grin.gov/gringlobal/accessiondetail?id=1600266; USDA Agriculture Research Service, 2015) The species distribution GIS information was downloaded from https://github.com/wpetry/USTreeAtlas/tree/main.

### DNA extractions

Frozen, dark-treated, fresh leaves from PI 613880 were used for DNA and high-molecular weight (HMW) extractions. For each extraction, 0.25 g of tissue were ground into a fine powder using liquid N_2_-chilled mortar and pestle. For Illumina sequencing, the DNA was obtained using a Plant Pro DNA extraction kit (Qiagen, Germantown, MD) with no additions to the manufacturer’s protocol. A Zymo Genomic DNA Clean & Concentration kit (Irvine, CA) was used to clean and concentrate the resulting gDNA samples. Sample quality and quantity was tested on a Nanodrop Spectrometer (Thermo Fisher Scientific, Waltham, MA) and Qubit 4 Fluorometer (Thermo Fisher Scientific, Waltham, MA). For HMW DNA extraction, we used a Qiagen Genomic-tip 100/G kit following the protocol developed by Driguez et al. (2021) with no modifications. HMW gDNA was checked for quality and quantity on a Nanodrop Spectrometer (Thermo Fisher Scientific, Waltham, MA) and Qubit 4 Fluorometer (Thermo Fisher Scientific, Waltham, MA). We further size selected HMW DNA samples using a Short Read Eliminator XL Kit (Circulomics, Baltimore, MD).

### Library Preparation and Sequencing

To obtain high-depth and high-quality short reads for PI 613880, we conducted whole genome sequencing (WGS) on the HiSeq platform (Illumina, San Diego, CA) in a paired-end format with a read length of 150 bp. GeneWiz (now Azenta, South Plainfield, NJ) performed the library preparations and sequencing as contractual service. We additionally had University of Maryland Institute of Genome Sciences (Baltimore, MD) prepare a PacBio HiFi library from the HMW DNA and sequence it on a Sequel II system. We prepared Hi-C libraries for PI 618880 using a Phase Genomic Proximo Plant Kit (Seattle, WA) and had contractual sequencing done by GeneWiz using a paired-end 150 bp read length format on a MiSeq (for library quality control testing) and HiSeq Illumina machines.

### RNA Extraction and Sequencing

Shoot apices were randomly collected from the living accession of PI 613880 at the USDA *Malus* Collection and flash frozen with liquid N_2_ and stored at −80°C. Collections for apices occurred on July 25th, August 28th, October 5th, and November 8th, 2018. For the July and October collection dates, each replicate sample was divided into three biological replicates of at least 25 apices/replicates. For the remaining three collection dates a single pooled sample of 5 apices/date was made. Each replicate apex sample was then ground into a fine powder using a mortar and pestle under liquid nitrogen, and RNA was extracted from the frozen ground tissue using a CTAB-based extraction method (Gasic et al. 2004). Spermidine was substituted for spermine in the extraction buffer. Extracted RNA was then further purified using a commercial kit (RNeasy Plant; Qiagen). Purified RNA was assessed for quality using 1.2% formaldehyde gel electrophoresis, and quantified by spectroscopy (NanoDrop 2000c, ThermoScientific, Waltham, MA). RNA libraries were prepared and sequenced by Novogene (Sacramento, CA) on the Illumina HiSeq platform generating paired-end, 150-base-pair length reads.

### Genome Assembly of PI 613880

To determine genome size and heterozygosity by *k*-mer counting using the WGS data, we first filtered the WGS for duplication using nubeam-dedup software (Dai & Guan, 2020). The filtered reads where then used as an input into KMC 3 (v3.0) (Kokot et al., 2017) using the parameters ‵-k21 -t10 -m64 -ci1 - cs1000000‵ to count *k-*mers. The resulting counts were then uploaded into GenomeScope (2.0) following the instructions (Ranallo-Benavidez et al., 2020), which provided an estimated genome size and heterozygosity. PacBio HiFi reads were filtered for adapters using HiFiAdapterFilt (v2.0.0; Sim et al., 2022). We then assembled a draft genome using the PacBio HiFi sequencing using the HiFiasm (v0.16.1) assembler with the Hi-C reads used as an additional input to facilitate phasing of the contigs without parental information (Cheng et al., 2022).

Our approach to scaffolding was replicated from Mansfeld et al. (2023). In short, the draft haplotype assemblies were scaffolded using Chromonomer (v.1.13) with linkage maps markers sourced from *M*. × *domestica* (Bianco et al., 2014; Di Pierro et al., 2016; Catchen et al., 2020). To validate the scaffolding and correct mis-joins, we mapped the Hi-C reads following processing using the Arima Pipeline (https://github.com/ArimaGenomics/mapping_pipeline) (Ghurye et al., 2017), Phase Genomics Juicebox utility Matlock (https://github.com/phasegenomics/matlock), and 3D-DNA run-assembly-visualizer script to generate a Juicebox editable assembly file (https://github.com/aidenlab/3d-dna/blob/master/visualize/run-assembly-visualizer.sh). Scaffolding corrections were manually performed using Juicebox (v1.11.08) (Durand et al., 2016).

The resulting scaffolded FASTA files for each haplotype were then aligned to GDDH13 v1.1 assembly using nucmer from the MUMmer package (v4.0.0beta2) (Marçais et al., 2018; Velasco et al., 2010). We used the mapping to assign chromosome numbers to each scaffold. However, Chromosome 05 required reverse-complementation. We also performed plastid sequence decontamination using the blobtools (v1.1.1) following the default protocol (Laetsch & Blaxter 2017). Genome statistics and quality were assessed using Busco (v.5.3) using the embryophyta ODB10 database, gt seqstat (v1.6.2), and Merqury (v1.4) (Rhie et al., 2020; Simão et al., 2015; Gremme et al., 2013).

### Haplotypic variation

Alignment of the two haplotype FASTA files was performed to identify structural variants (SVs) using the nucmer tool (Marçais et al., 2018) with specific parameters ‵--maxmatch -l 100 -c 500‵. First, SNPs and Indels were identified using nucmer ‵show-snps -CT‵. Additionally, the delta file was submitted to the Assemblytics web interface (http://www.assemblytics.com/) for analysis, where a maximum variant size of 50 kbp was defined according to Nattestad & Schatz (2016). The analysis results were then retrieved in the form of a BED file and imported into R for visualization of the size and location distributions.

### Genome Annotation

The annotation of long-terminal repeats (LTR) was performed with EDTA (v1.9.6) (Ou et al., 2019) with species ‵-others‵ option enabled and all other parameters set to default. We assessed assembly quality using the LAI program (Ou et al., 2018) within the LTR_retriever software (v2.9.0) (Oh & Jiang, 2018). For annotation of the genes, we employed a comprehensive approach using the MAKER2 (v2.31.1) pipeline (Holt and Yandell 2011). First, the 7 RNA-seq libraries were evaluated for adapter contamination and quality using FASTQC (v.0.11.8) (https://www.bioinformatics.babraham.ac.uk/projects/fastqc/). The libraries were then mapped to each chromosome-only haplotype assembly using STAR aligner with the parameters ‵--outSAMstrandField intronMotif‵ and ‵--alignIntronMaxenabled 10 kb‵ (Dobin et al., 2013).

Transcripts for each haplotype were then assembled independently using the read alignments as input into the transcript assembler StringTie2 (v2.2.1) (Kovaka et al., 2019). GFFread (v0.12.7) was used to generate a FASTA file for the StringTie2 transcripts (Pertea and Pertea 2020). We then used the StringTie2 assembled transcripts as pseudo-expressed sequence tags (EST) evidence in the first round of evidence-based annotation using MAKER2. The first round of annotation was then extracted using the extract_anno_evi.sh script and used as training data for *ab initio* gene predictors SNAP and Augustus (Korf 2004; Hoff & Stanke 2019). We followed the first round of *ab initio* gene prediction with a second round of MAKER2 using the optimized transcripts from the first round as models. Following the second round of *ab initio* gene prediction, the optimized transcripts were used as input into a final round of MAKER2. We additionally annotated the rRNA and tRNA features using Infernal (v1.1.4) and tRNAscan-SE (v2.0) (Lagesen et al., 2007; Chan & Lowe 2019). Gene annotation stats and extraction of transcript and translated peptide sequences were performed using AGAT (v0.9.1) (Dainet 2019). The distribution of genomic features and structural variation was plotted by importing the respective GFFand BED files into R. Genomic regions were binned into 100 kb regions and the relative size of each feature within each bin. The respective values were then plotted as a heatmap in ggplot2.

### BUSCO based phylogeny

For each Malus species genome analyzed, we ran a BUSCO analysis as above. Single copy BUSCO genes were retrieved concatenated and aligned using the BUSCO_phylogenomics python script https://github.com/jamiemcg/BUSCO_phylogenomics. The multisequence alignment fasta file was then imported into IQ-tree2 (Nguyen et al., 2015) using the ModelFinder module (Kalyaanamoorthy et al., 2017) to identify the best substitution model. We used the ultrafast bootstrap approximation with 1000 replicates (Hoang et al., 2017) to establish branch support values. The maximum likelihood tree was then plotted in the interactive tree of life web interface (Letunic and Bork 2021).

### Synteny analyses

A pairwise comparison of gene synteny between the wild *Malus* and *M.* × *domestica* genomes published genomes. The pairwise comparison order was determined based on the phylogenetic analyses outlined above. The comparison sequence involved the *M. angustifolia* haplotype 1 assembly being compared to *M. fusca* haplotype 1 (Mansfeld et al., 2023), followed by a comparison of *M. fusca* to *M. sieversii* haploid, then *M. sieversii* to *M. sylvestris* haploid (Sun et al., 2020), and finally *M. sylvestris* to the *M.* × *domestica* genome cv. ‘Honeycrisp’ haplotype 1 (Khan et al., 2022). Synteny was determined using the Python MCScanX pipeline v1.1.12 (Tang et al., 2008; Wang et al., 2012), and by aligning the coding sequences of the gene annotations for each species pair. For the purpose of plotting karyotypes, macro level synteny was defined with a minimum syntenic block size of 10 genes (‵--minspan 10‵). Additionally, an *M. angustifolia* vs. all analysis was also performed using the same methods.

### Data and Script Availability

The raw sequence data generated from this project can be retrieved from NCBI SRA database upon publication to a journal. Final assemblies for both haplotype genomes can be retrieved from the Genome Database for Rosaceae (GDR) (Jung et al., 2019). Scripts for publishing the figures will be made available at github.com/bmansfeld/mangustifolia_figs.

## Results

### Assembly of a chromosomal-scale haplotype resolved genome

We obtained 64.4 Gb of Illumina sequence data, which was used to count *k-*mer frequency. Using those *k-*mers counts, GenomeScope estimated *M. angustifolia* PI 613880 to have a genome size of 747 Mb and was expected to be diploid with a heterozygosity of 1.06% (Fig 1C). We also generated 1.8 million HiFi reads which totaled 22.4 Gb of sequence. The HiFi sequencing is estimated to be ∼30x of the estimated genome size, which was of sufficient depth to conduct *de novo* genome assembly using HiFiasm (Cheng et al., 2021, Cheng et al., 2022). Additionally, we obtained 57.5 Gb of Hi-C data to be used during the phasing and scaffolding steps. Using all of the available sequence data, we were able to yield a final assembly for two phased haplotypes of ∼717 Mb (Table 1). Each assembly was contained in less than 400 contigs/scaffolds and exhibited 17 chromosome-scale pseudomolecules after scaffolding (Fig 2A and C). Our N50 value for each haplotype was greater than 40.7 Mb and exhibited a QV value of 63. Sub-setting the assemblies only for the chromosome-scale molecules (*i.e.,* removing additional un-scaffolded contigs and sequencing artifacts) yielded a reduction in assembly length of <12 Mb and had no effect on N50. BUSCO assessment indicated highly complete chromosome-only assemblies with greater than 98.7% complete BUSCO genes of the 1614 contained in the Embryophyta OBD10 database (Table 1).

**Figure 2.**
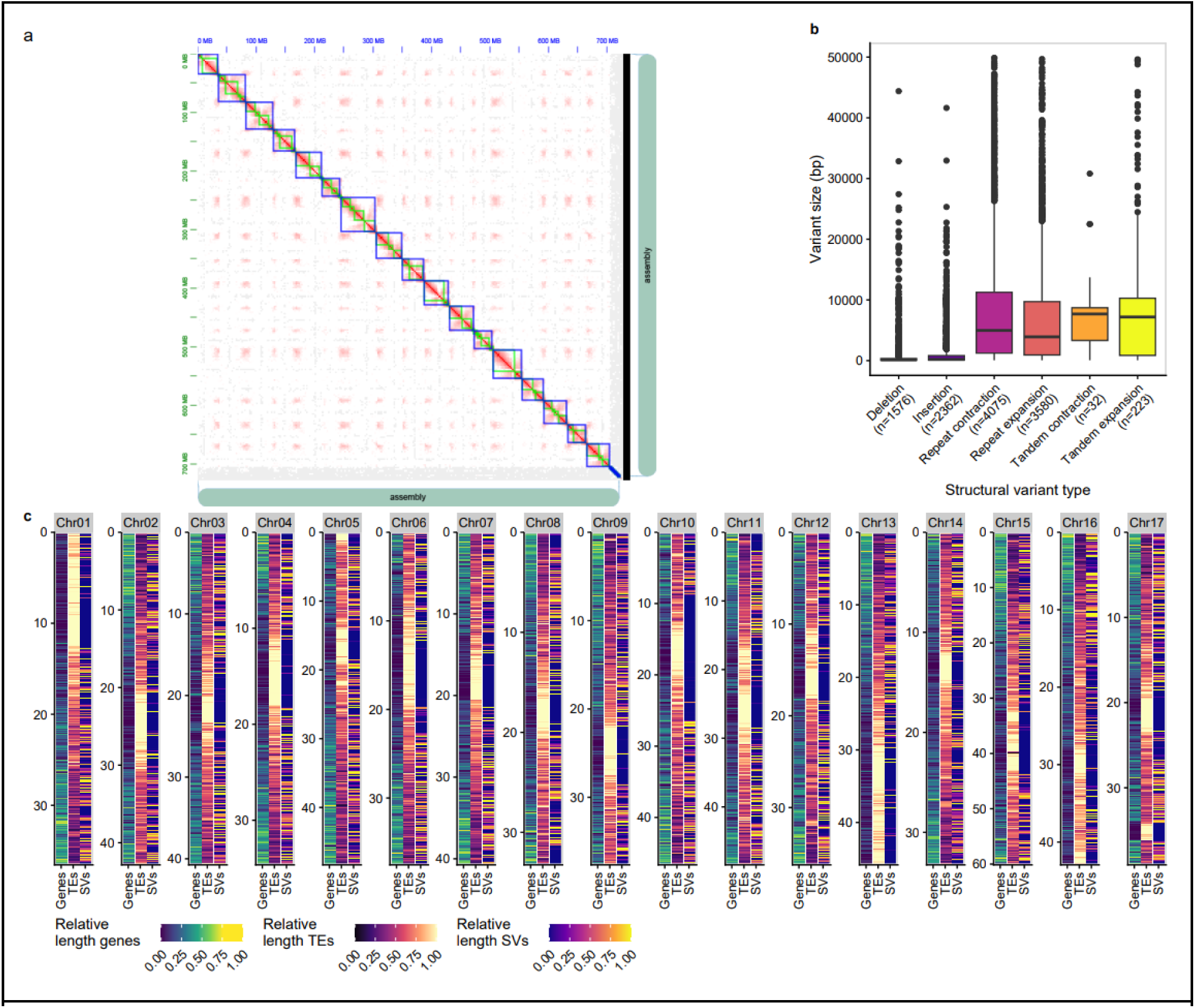
Genome assembly of *Malus angustifolia.* A) Hi-C contact map of Haplotype 1. B) Hi-C contact map of Haplotype 1. C) Heat maps of the annotated features distributed across all 17 assembled chromosomes (genes, TEs-Transposable Elements, and SVs-Structural Variations). A 100 Kbp window was used to calculate the relative cumulative length of each feature type. D) Comparison of haplotypic SV distributed by length.

**Table 1.** Genome assembly statistics.

| Assembly | Haplotype | Contig/Scaffold Number | Assembly Length | Contigs >1 Mb | N50 | QV | BUSCO<sup>1</sup> |
| --- | --- | --- | --- | --- | --- | --- | --- |
| Scaffolded Assembly | Hap1 | 396 | 717,543,921 | 17 | 40,768,280 | 63.04 | - |
|  | Hap2 | 331 | 717,616,600 | 17 | 40,956,496 | 63.04 | - |
| Chromosome-only | Hap1 | 17 | 705,089,958 | 17 | 40,768,280 | - | C:98.8%[S:62.1%,D:36.7%],F:0.6%,M:0.6% |
|  | Hap2 | 17 | 703,144,148 | 17 | 40,956,496 | - | C:98.7%[S:62.3%,D:36.4%],F:0.6%,M:0.7% |

<sup>1</sup>BUSCO - Benchmark Universal Single Copy Orthologs; C - Complete BUSCOs; S - Complete and single-copy BUSCOs; D - Complete and Duplicated BUSCOs; F - Fragmented BUSCOs; M - Missing BUSCO; n = 1614

### Annotation of M. angustifolia haplotype assemblies

To annotate the long-terminal repeat (LTR) retrotransposon space, we employed a pan-genome approach within the EDTA software package (Ou et al., 2019). This approach was successfully used to annotate the *M. fusca* v1.1 genome (Mansfeld et al., 2023). We identified that each *M. angustifolia* haplotype assembly contained 58.6% LTRs (Table 2), roughly 4% more than the smaller genomes of its other wild relatives. Compared to the *M. fusca* genome, *M. angustifolia* contained 41.75 - 47.38 Mb more bp associated with annotated repeats. Unknown LTRs features contributed the most to the increased amount of repeats in *M. angustifolia.* However, the percent of the *M. angustifolia* genome that was annotated as repeats was ∼58%, which is similar to the percent reported for *M. fusca* but lower than the ∼62% reported for ‘Honeycrisp’ (Khan et al., 2022; Mansfeld et al., 2023). We further used LTR annotation to assess the assembly quality and contiguity of repeat space (i.e., by using the LTR Assembly Index (LAI)) to assess quality in assembling the LTR space, for which each haplotype was greater than 21 (21.88 and 22.07, respectively) (Ou et al., 2018). LAI values are the highest report for a *Malus* assembly and correspond to a “Gold Standard” (Mansfeld et al., 2023; Ou et al., 2018).

**Table 2.** Genome annotation statistics for the chromosome-only genome assembly.

| Assembly | Haplotype 1 | Haplotype 2 | Combined |
| --- | --- | --- | --- |
| *Intergenic* |  |  |  |
| LTR - Copia | 10.57% | 10.74% |  |
| LTR - Gypsy | 15.75% | 15.65% |  |
| LTR - Unknown | 21.56% | 21.44% |  |
| TIR-CACTA | 1.26% | 1.28% |  |
| TIR-Mutator | 4.27% | 4.23% |  |
| TIR-PIF Harbinger | 2.41% | 2.40% |  |
| TIR-Tc1 Mariner | 0.03% | 0.03% |  |
| TIR-hAT | 2.50% | 2.57% |  |
| Low complexity | 0.02% | 0.02% |  |
| nonTIR - helitron | 0.26% | 0.29% |  |
| Total % | 58.63% | 58.64% |  |
| Total bp | 413,396,256 | 412,293,630 |  |
| LAI | 22.07 | 21.88 |  |
| *Genic* |  |  |  |
| % Complete BUSCO genes | 98.1 | 98.7 |  |
| Complete single copy BUSCO genes | 1003 | 1006 |  |
| Complete duplicate BUSCO genes | 592 | 587 |  |
| Fragmented BUSCO genes | 9 | 10 |  |
| Missing BUSCO genes | 10 | 11 |  |
| Genes | 43,949 | 47,334 | 91,283 |
| tRNA genes | 717 | 934 | 1651 |
| rRNA gene | 450 | 440 | 890 |

Footnote: BUSCO (v.5.3) using the embrophyta odb 10, $n=1614$

To annotate the gene space, we utilized a hybrid approach that mixed transcriptional evidence and *ab initio* prediction within the MAKER2 suite (Holt and Yandell 2011). To do this, RNA sequencing was performed on RNA extract from shoot apices and generated a total of ∼55.4 Gbp of sequence, with an average of ∼26.4 million reads per library (STable 1). We mapped over 184 M reads to each haplotype genome using STAR aligner, assembled pseudo-EST evidence with StringTie2, and trained gene prediction software based on those transcript models within the MAKER2 pipeline (Holt & Yandell, 2011; Dobin et al., 2013; Kovaka et al., 2019). Of the two haplotype assemblies, Hap2 had ∼47,000 genes annotated and had a BUSCO Complete score of 95.9%. Whereas, Hap1 had ∼44,000 genes annotated and had a BUSCO Complete score of 96.0%. We recently annotated the *M. fusca* genome using similar approaches but leveraging a substantial volume of RNAseq data from highly varied libraries. Those annotations only yielded marginally higher BUSCO transcriptome scores (*i.e.,* 96.8% compete for *M. fusca* haplotype 1), indicating that while a more limited data set was used herein, it was sufficient to accurately represent the annotation for *M. angustifolia*. We additionally annotated the rRNA and tRNA space and found each haplotype contained 440-450 rRNA and 717-934 tRNA genes (Table 2).

### Haplotypic variation in M. angustifolia is largely at the single nucleotide level

We conducted comparative haplotypic analysis by aligning the two haplotype assemblies and detected SNPs and substantial structural variations (SVs) between them. Our findings revealed a total of 2,941,850 heterozygous SNPs, approximately 500,000 more than that observed in the closely related *M. fusca* genome. Additionally, 1,977,328 small InDels were identified between the haplotypes - roughly the same amount as in *M. fusca*. Notably, in plants with high heterozygosity, the source of genetic diversity extends beyond single nucleotide polymorphisms, and it has been shown that large structural variations (SVs) play a significant role in haplotypic diversity (Guan et al., 2021; Mansfeld et al., 2021; Yuan et al., 2021; Massonnet et al., 2022). Using our approach, we identified SVs of different types ranging in size from 50 bp - 50 Kb. Interestingly, while the heterozygosity rate of *M. angustifolia* (∼1.1%) is higher than that described in *M. fusca* (∼0.8%), we identified about half of the SVs in *M. angustifolia* compared to *M. fusca*. For example, while 3,774 large haplotypic deletions were observed in *M. fusca*, only ∼1500 were observed in *M. angustifolia* (Fig 2B). This result suggests that, at least for the accession selected, heterozygosity levels are largely driven by the increase in SNPs rather than large SVs. It will be interesting to explore how population dynamics in the different species have impacted accumulation of large SVs, which are often considered to be deleterious (Zhou et al., 2019).

### Recapitulating the malus phylogeny and exploring synteny within wild Malus genomes

The assembly and annotation of the *M. angustifolia* genome in this study, along with our recent assembly of *M. fusca*, marks a significant milestone in accumulating a substantial repertoire of wild *Malus* assemblies. These resources enable, for the first time, a detailed comparative analysis between Eurasian (*M. sieversii* and *M. sylvestris*) and North American wild *Malus* species (*M. angustifolia* and *M. fusca*). While a phylogenetic framework for *Malus* x *domestica* and some of its wild relatives exists, it is largely based on plastid genomes (Nikiforova et al., 2013; Volk et al., 2015b) morphological traits, and/or based on specifically classified families of genes (Robinson et al., 2000; Forte et al., 2002). We sought to further validate the relationships between the sequenced wild species on a whole genome scale using recently published high-quality, chromosomal-scale assemblies (Sun et al., 2020; Khan et al., 2022; Mansfeld et al., 2023). To this end, we identified and aligned all single copy BUSCO genes in the *M. angustifolia* assembly and the other four species, namely *M. fusca, M. seiversii, M. sylvestris,* and *M.* x *domestica* ‘Honeycrisp’. By comparing single-copy BUSCO genes, we were able to construct a robust and well-supported maximum likelihood phylogenetic tree (Fig 3a), which highlighted the divergence and shared ancestry among these closely related *Malus* species. This served as a foundational step in validating previously hypothesized evolutionary dynamics, and taxonomic relationships within the genus.

**Figure 3.**
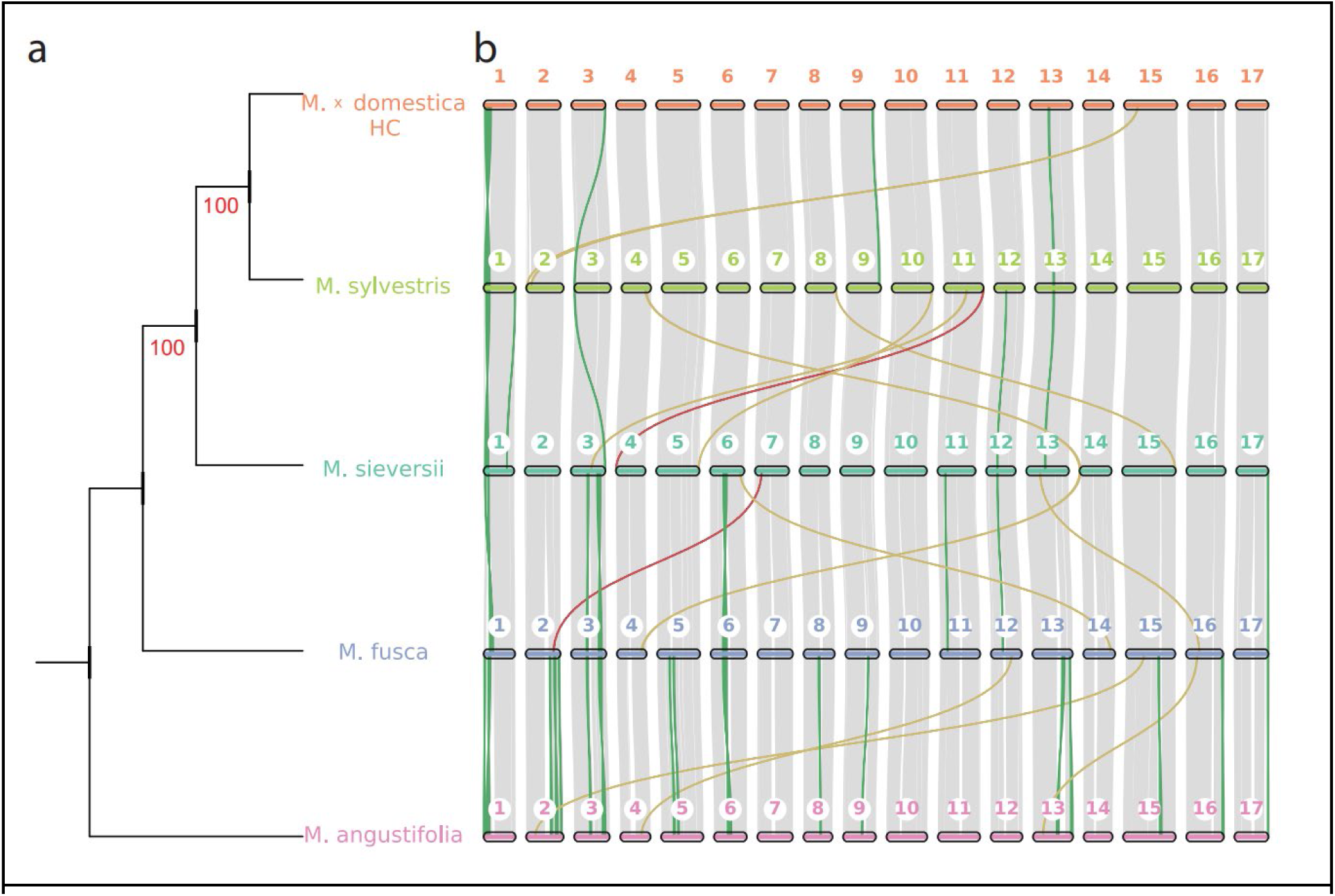
A) Phylogenetic relationships between Eurasian and North American Malus species. The maximum likelihood bootstrapped (n=1000) tree was constructed by aligning single copy BUSCO genes identified from each species using IQ-tree2. B) Synteny and collinearity within *Malus* crop wild relatives and representative *M. domestica* ‘Honeycrisp’. Green, yellow and red ribbons indicate inversions, translocations and inverted translocations, respectively.

We then leveraged these re-established phylogenetic relationships as a foundation and conducted a comprehensive pairwise synteny analysis between Eurasian and North American Malus species. Our analysis unveiled extensive synteny and conserved genomic segments across these species; however, it also underscored the accumulation of macro-level structural variations as one moves further away from *M.* x *domestica*. Notably, minimal structural divergence was observed between *M.* x *domestica* and its Eurasian progenitors, in contrast to the increasing variations apparent in the genomes of their North American counterparts. Notably, *M. angustifolia* genome emerged as the most distinct, displaying numerous inversions and translocations in comparison to the other species (Fig 3b).

This investigation enhances our understanding of the genomic architecture within the *Malus* genus and contributes to the elucidation of evolutionary dynamics and the genetic underpinnings of valuable agronomic traits in these wild apple species. Considering the extensive insights gained from this study, there is a compelling case for further endeavors to assemble additional wild *Malus* genomes. Expanding the repertoire of sequenced wild *Malus* genomes promises to provide an even more comprehensive understanding of the genus’s evolutionary history, genomic diversity, and the genetic reservoir that underpins valuable agricultural traits, ultimately facilitating more effective conservation and breeding strategies for these important fruit trees.

## Supplemental Tables

**STable 1.** RNA samples used in the annotation of the *M. angustifolia* PI 613880 genome.

| Collection Date | Biological Replicate Number | Number of Reads |
| --- | --- | --- |
| 7/25/2018 | 1 | 27,162,705 |
| 7/25/2018 | 2 | 28,401,252 |
| 7/25/2018 | 3 | 25,529,522 |
| 10/5/2018 | 1 | 29,705,576 |
| 10/5/2018 | 2 | 23,512,852 |
| 10/5/2018 | 3 | 22,873,449 |
| 6/1/2018, 8/28/2018, 11/8/2018 | 1 | 27,388,712 |
| **Total** |  | **184,574,068** |

